# Compositional profiling of EV-lipoprotein mixtures by AFM nanomechanical imaging

**DOI:** 10.1101/2022.07.19.500441

**Authors:** Andrea Ridolfi, Laura Conti, Marco Brucale, Roberto Frigerio, Jacopo Cardellini, Angelo Musicò, Miriam Romano, Andrea Zendrini, Laura Polito, Greta Bergamaschi, Alessandro Gori, Costanza Montis, Lucio Barile, Debora Berti, Annalisa Radeghieri, Paolo Bergese, Marina Cretich, Francesco Valle

**Affiliations:** Consiglio Nazionale delle Ricerche, Istituto per lo Studio dei Materiali Nanostrutturati, Bologna, Italy; Dipartimento di Chimica “Ugo Schiff”, Università degli Studi di Firenze, Firenze, Italy; Consorzio Interuniversitario per lo Sviluppo dei Sistemi a Grande Interfase, Firenze, Italy; Consiglio Nazionale delle Ricerche, Istituto di Scienze e Tecnologie Chimiche “Giulio Natta”, Milan, Italy; Dipartimento di Medicina Molecolare e Traslazionale, Università degli Studi di Brescia, Brescia, Italy; Istituto Cardiocentro Ticino, Ente Ospedaliero Cantonale, Lugano, Switzerland; Consiglio Nazionale delle Ricerche, Istituto per la Ricerca e l’innovazione Biomedica, Palermo, Italy

## Abstract

The widely overlapping physicochemical properties of lipoproteins (LPs) and extracellular vesicles (EVs) represents one of the main obstacles for the isolation and characterization of these pervasive biogenic lipid nanoparticles. We herein present the application of an atomic force microscopy (AFM)-based quantitative morphometry assay to the rapid nanomechanical screening of mixed LPs and EVs samples.

The method can determine the diameter and the mechanical stiffness of hundreds of individual nanometric objects within few hours. The obtained diameters are in quantitative accord with those measured via cryo-electron microscopy (cryo-EM); the assignment of a specific nanomechanical readout to each object enables the simultaneous discrimination of co-isolated EVs and LPs even if they have overlapping size distributions. EVs and all classes of LPs are shown to be characterized by specific combinations of diameter and stiffness, thus making it possible to estimate their relative abundance in EV/LP mixed samples in terms of stoichiometric ratio, surface area and volume. As a side finding, we show how the mechanical behaviour of specific LP classes is correlated to distinctive structural features revealed by cryo-EM. To the best of our knowledge, these results represent the first systematic single-particle mechanical investigation of lipoproteins.

The described approach is label-free, single-step and relatively quick to perform. Importantly, it can be used to analyze samples which prove very challenging to assess with several established techniques due to ensemble-averaging, low sensibility to small particles, or both, thus providing a very useful tool for quickly assessing the purity of EV/LP isolates including plasma- and serum-derived preparations.

## Introduction

Extracellular vesicles (EVs) are membranous nanoparticles released by cells as mediators of physiological and pathological processes. They are able to shuttle nucleic acids, proteins and lipids to distant targets and are considered key players of intercellular communication [Maas 2017].

Lipoproteins (LPs) are a class of nanobioparticles pervasively found in interstitial fluid [Sloop 1987, Busatto 2020], plasma and serum [Geeurickx 2020; Freitas 2019], as well as in conditioned culture medium [Zhang 2020] and milk [Hu 2021]. Their primary function is the dispersion of lipids to facilitate their transport and delivery [Ramasamy 2014].

LPs have sizes, densities, and surface compositions overlapping those of extracellular vesicles (EVs) [Simonsen 2017; Busatto 2019]; moreover, in most sources the relative abundance of LPs is several orders of magnitude higher than EVs in terms of number density [Johnsen 2019]. Due to this, detecting and separating LPs in EV preparations is very challenging [Botha 2022]. As a consequence, EVs and LPs are often co-isolated from several sources, notably including plasma and serum [Holcar 2021; Brennan 2020; De Rond 2019; van der Pol 2018; Sodar 2016]. The detection and quantification of LPs in complex samples are thus key issues for the overall reproducibility of EV research [Nieuwland 2022] and for the study of EV/LP functional interplay [Busatto 2022; Busatto 2020]. However, many of the most widespread nanoparticle characterization techniques fail to deal with the extremely wide range of sizes and densities encompassed by different LPs classes; due to this, LP assessment remains problematic [Mørk 2017; Nieuwland 2022].

Recently, atomic force microscopy (AFM)-based nanomechanics have been established as a valuable tool to assess EV identity, purity and function [LeClaire 2021]: EVs have been shown to display a specific nanomechanical fingerprint which is detectable via force spectroscopy [Vorselen 2017; Piontek 2021] and which can be leveraged to discriminate between different EV populations [Sorkin 2018; Vorselen 2018]. We recently implemented a single-particle nanomechanical screening based on quantitative AFM morphometry which considerably increases analytical throughput [Ridolfi 2020a], making it possible to rapidly detect co-isolated, non-vesicular contaminants [Ridolfi 2020a; Ridolfi 2020b] and to give an estimate of their abundance relative to that of EVs [Borup 2022]. Conc4versely, the nanomechanical behaviour of LPs is still largely uncharacterized, the first studies having appeared only very recently [Baraimukov 2022].

We first formulated the working hypothesis that the same method we developed for EVs [Ridolfi 2020a] might be applicable to LPs and thus be able to differentiate between LP subtypes on a nanomechanical basis. To test the validity of this hypothesis, we applied the AFM method mentioned above to a set of commercially isolated LPs and to two models of human EVs with different size distributions and biogenesis pathways: human Cardiac Progenitor Cell EVs (hCPC-EVs) [Andriolo 2018] and human Red Blood cell derived EVs (RBC-EVs) [Usman 2018].

We find that our method is able to discern the specific nanomechanical fingerprint of individual LP subclasses and EVs, thus enabling the detection, quantification and size distribution determination of specific subpopulations in complex EV/LP mixtures. As a proof of concept, we show that our method can perform the single-step, label-free determination of an ultracentrifuged plasma sample, determining the size distributions of individual LP/EV subtypes in the mixture and estimating their relative abundances. As a side finding, we note how the mechanical behaviour of specific LP classes is correlated to distinctive structural features as detected by cryo-EM. To the best of our knowledge, these results represent the first systematic single-particle mechanical investigation of lipoproteins.

## Materials and Methods

### EVs and LPs samples preparation

Low Density (LDL), High Density (HDL) and Very Low Density (VLDL) lipoproteins were acquired from MyBioSource (San Diego, CA); Intermediate Density Lipoprotein (IDL) were purchased from LSBio (Seattle, WA), Chylomicrons were purchased from BioVision (Waltham, MA). Red blood cells EVs (RBC-EVs) were separated from healthy donors’ red blood cells (see Supporting Information) as described elsewhere [Usman 2018]. Human Cardiac Progenitor Cell EVs (hCPC-EVs) were isolated as described elsewhere [Andriolo 2018]. Platelet-free plasma from healthy donors was purchased from Cerba Xpert (Saint Ouen L’Aumone, France) and centrifuged at 100k x g for 120 min; after removing the supernatant, the resulting pellet was resuspended in 200 µl of PBS. Ultrapure water was prepared with a Millipore Simplicity UV apparatus. All other reagents were acquired from Sigma-Aldrich Inc (www.sigmaaldrich.com) unless otherwise stated.

### BCA Assay

Protein concentrations in lipoprotein and EV samples were determined with Pierce(tm) BCA Protein Assay Kit (ThermoFisher, Rockford, USA) according to manufacturer instructions. 25 µl of samples or BSA standards were pipetted into the microplate wells; 200 µl of working reagent was added in the plate and incubated at 37 °C for 30 minutes. The plate was read at 560 nm on the plate reader (HiPo MPP-96 Microplate Photometer, Biosan, Riga, LV).

### Lipid Assay

Lipid concentrations were determined with a Lipid Quantification Kit (Cell Biolabs, Inc). 15 µl of analyte or of a DOPC standard were added to 150 µl of 18 M H_2_SO_4_ and successively incubated at 90°C for 10’ and at 4 °C for 5’. 100 µl of the mixture were transferred into a 96-well plate and additioned with 100 µl of the sulfo-phopsho-vanillin reagent. The plate was incubated for 15 minutes a t 37 °C and read with the plate reader (HiPo MPP-96 Microplate Photometer, Biosan, Riga, LV) at 520 nm wavelength.

### Western Blotting

8 µl of (5x) Laemmli buffer were added to 32 µl of LPs/EVs solutions. Samples were then heated for 10’ at 95 °C. Proteins were separated by SDS-PAGE (4– 20%, Mini-Protean TGX Precast protein gel, Bio-Rad) and transferred onto a nitrocellulose membrane (BioRad, Trans-Blot Turbo). Nonspecific sites were saturated with a TBS-T solution (0,05% Tween-20) with 1% BSA for 1 h. For RBC-EV samples, membranes were incubated overnight at 4°C with: mouse anti-GM130 (1:1000 BD Biosciences, Germany), mouse anti-Alix (1:1000 Santa Cruz Biotechnology, USA), rabbit ant-Annexin XI (1:1000 GeneTex, USA), mouse anti-LAMP1 (1:1000 Santa Cruz Biotechnology, USA), anti-HBB (1:1000 Abnova, Jhouzih St., Taipei, Taiwan). For hCPC-EV samples, membranes were incubated overnight at 4 °C with anti-CD9 (1:1000, BD Pharmingen), anti-CD63 (1:1000; BD Pharmingen, San Jose, CA, USA), anti-Alix (1:1000, Santa Cruz, CA, USA), and anti-TSG101 (1:1000, Novus Bio, Centennial, CO, USA). For LPs samples, membranes were incubated with anti-ApoA1 (1:1000, Santa Cruz, CA, USA), anti-ApoE (1:500, Santa Cruz, CA, USA) and anti-ApoB (1:500, Santa Cruz, CA, USA). After washing with TBS-T, membranes were incubated with horseradish peroxidase-conjugated (Jackson ImmunoResearch, Tucker, GA, USA) secondary antibodies diluted 1:5000 in TBS-T with 1% BSA for 1 h. After rinsing, the signal was developed using Bio-Rad Clarity Western ECL Substrate (Bio-Rad) and imaged using a Chemidoc XRS+ (BioRad).

### Nanoparticle Tracking Analysis

Nanoparticle tracking analysis (NTA) was performed according to the manufacturer’s instructions using a NanoSight NS300 system (Malvern Technologies, Malvern, UK) configured with a 532 nm laser. Samples were diluted in micro-filtered PBS; the ideal measurement concentrations were identified by pre-testing the ideal particle per frame value (20–100 particles/frame). A syringe pump with constant flow injection was used and three videos of 60 s were captured and analyzed with Malvern NTA software version 3.4. From each video, the mean, mode, and median EVs size was used to calculate sample concentration, expressed as nanoparticles/ml. It was not possible to analyse HDL, LDL and IDL samples because the particles diameter is below the detection range of the instrument.

### Dynamic Light Scattering (DLS) and ζ-potential

Samples were diluted in micro-filtered (0.22 µm) PBS to a final volume of 3 ml. Dynamic light scattering (DLS) was performed on a 90Plus particle size analyzer from Brookhaven Instrument Corporation (Holtsville, NY, USA) operating at 15 mW of a solid-state laser (λ = 661 nm), using a scattering angle of 90°. Each sample was equilibrated at 25 °C for 3’ prior to measurement. Mie theory was used to calculate the hydrodynamic diameter (Hd), considering absolute viscosity and refractive index values of the medium to be 0.890 cP and 1.330, respectively. The ζ-potential was determined at 25 °C using the same instrument equipped with an AQ-809 electrode, operating at applied current 150mA. The ζ-potential was calculated from electrophoretic mobility based on the Smoluchowski theory, assuming a viscosity of 0.890 cP and a dielectric constant of 78.5.

### Cryo-Electron Microscopy

3 μl of each sample were applied on glow-discharged Quantifoil Cu 300 R2/2 grids, then plunge-frozen in liquid ethane using a FEI Vitrobot Mark IV (Thermo Fisher Scientific) instrument. Excess liquid was removed by blotting for 1 s (blot force of 1) using filter paper under 100% humidity at 10 °C. Cryo-EM data were collected at the Florence Center for Electron Nanoscopy (FloCEN), University of Florence, on a Glacios (Thermo Fisher Scientific) instrument at 200 kV equipped with a Falcon III detector operated in the counting mode. Images were acquired using EPU software with a physical pixel size of 2.5 Å and a total electron dose of ∼ 50 e^-^/Å^2^ per micrograph. The diameters of individual EVs and LPs were estimated by averaging the minimum and maximum Feret diameter of their projection via Fiji [Schindelin 2012]. Measurements from several individual objects were pooled (N between 42 and 106 for different samples) to reconstruct diameter distributions.

### Atomic Force Microscopy

AFM imaging was performed on poly-L-lysine (PLL)-coated glass coverslips prepared following a revised version of the protocol described in [Ridolfi 2020a] which optimizes reproducibility. Microscopy glass slides (15mm diameter round coverslips, Menzel Gläser) were first incubated for 2 h in a 3:1 (v:v) 96% H_2_SO_4_/30% H_2_O_2_ ‘piranha’ solution, rinsed extensively in ultrapure water, cleaned in a sonicator bath (Elmasonic Elma S30H) for 30’ in acetone, followed by 30’ in isopropanol and 30’ in ultrapure water, and finally activated with air plasma (Pelco EasiGlow) for 5’ followed by immediate immersion in ultrapure water. Clean slides were then incubated for 30’ in a 0.01 (mg/ml) freshly prepared PLL solution in 100 mM, pH 8.5 borate buffer at room temperature, thoroughly rinsed with ultrapure water and dried with a gentle nitrogen flow. Following this protocol, the water contact angle (1μl droplets at ∼25°C, measured with a GBX DigiDrop goniometer) of functionalized slides was 35°±3°.

A 10 μl droplet of the sample was deposited on a PLL-functionalized glass slide and left to adsorb for 30 minutes at 4°C, then inserted in the AFM fluid cell (see below) without further rinsing. The concentration of each sample was adjusted by trial and error in successive depositions in order to maximize the surface density of isolated, individual objects. Some of the commercial LP samples (e.g. HDL) needed to be diluted up to 10^6^ times to avoid the formation of clusters of adjoining objects.

All AFM experiments were performed in ultrapure water at room temperature on a Bruker Multimode8 equipped with Nanoscope V electronics, a sealed fluid cell and a type JV piezoelectric scanner using Bruker ScanAsystFluid+ probes (triangular cantilever, nominal tip curvature radius 2-12 nm, nominal elastic constant 0.7 N/m) calibrated with the thermal noise method [Hutter 1993].

Imaging was performed in PeakForce mode as described elsewhere [Ridolfi 2020a]. Image background subtraction was performed using Gwyddion 2.58 [Nečas 2012]. Image analysis was performed with a combination of Gwyddion and custom Python scripts to recover the surface contact angle and equivalent solution diameter of individual objects [Ridolfi 2020a]. Equivalent solution diameter (D_eq_) and equivalent spherical cap contact angle (CA) distributions were reconstructed by pooling the AFM morphometry measurements of several individual objects (N between 135 and 510 for different samples).

## Results and Discussion

### Characterizations of isolated LPs and EVs

Commercial purified LP samples (HDLs, IDLs, LDLs, VLDLs and Chylomicrons), together with hCPC-EVs and RBC-EVs, were first characterized by different techniques including Nanoparticle Tracking Analysis (NTA), Dynamic Light Scattering (DLS), ζ-potential determination, and Western Blot. For all LP samples, protein and lipid contents were measured by BCA and sulfo-phospho-vanillin assays. All the results of these characterizations were in line with expected values or outcomes for each LP subclass, including e.g. their average size, ζ-potential, protein/lipid ratio and the immunodetection of LP-associated apolipoproteins. Analyses of both hCPC-EVs and RBC-EVs were in agreement with previously published data [Andriolo 2018, Usman 2018]. Full details of the characterizations are reported in the Supporting Information.

### Ultrastructure of LPs and EVs via cryo-EM

We then collected Cryo-EM micrographs of each purified LP sample and of hCPC-EVs (see materials and methods), finding that recurring ultrastructural details were associated to specific samples (Figures 1 and 2).

**Figure 1.**
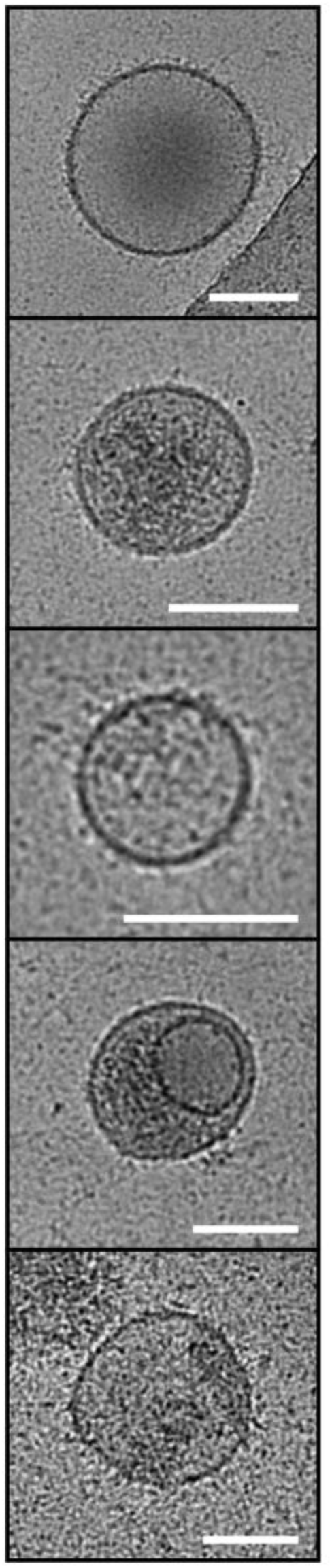
Representative cryo-EM images of individual hCPC-EVs. All scalebars are 100 nm. Several recurring structural features are visible: a high-contrast boundary corresponding to the bilayer, extensive external decoration, and occasional hints of luminal cargo.

**Figure 2.**
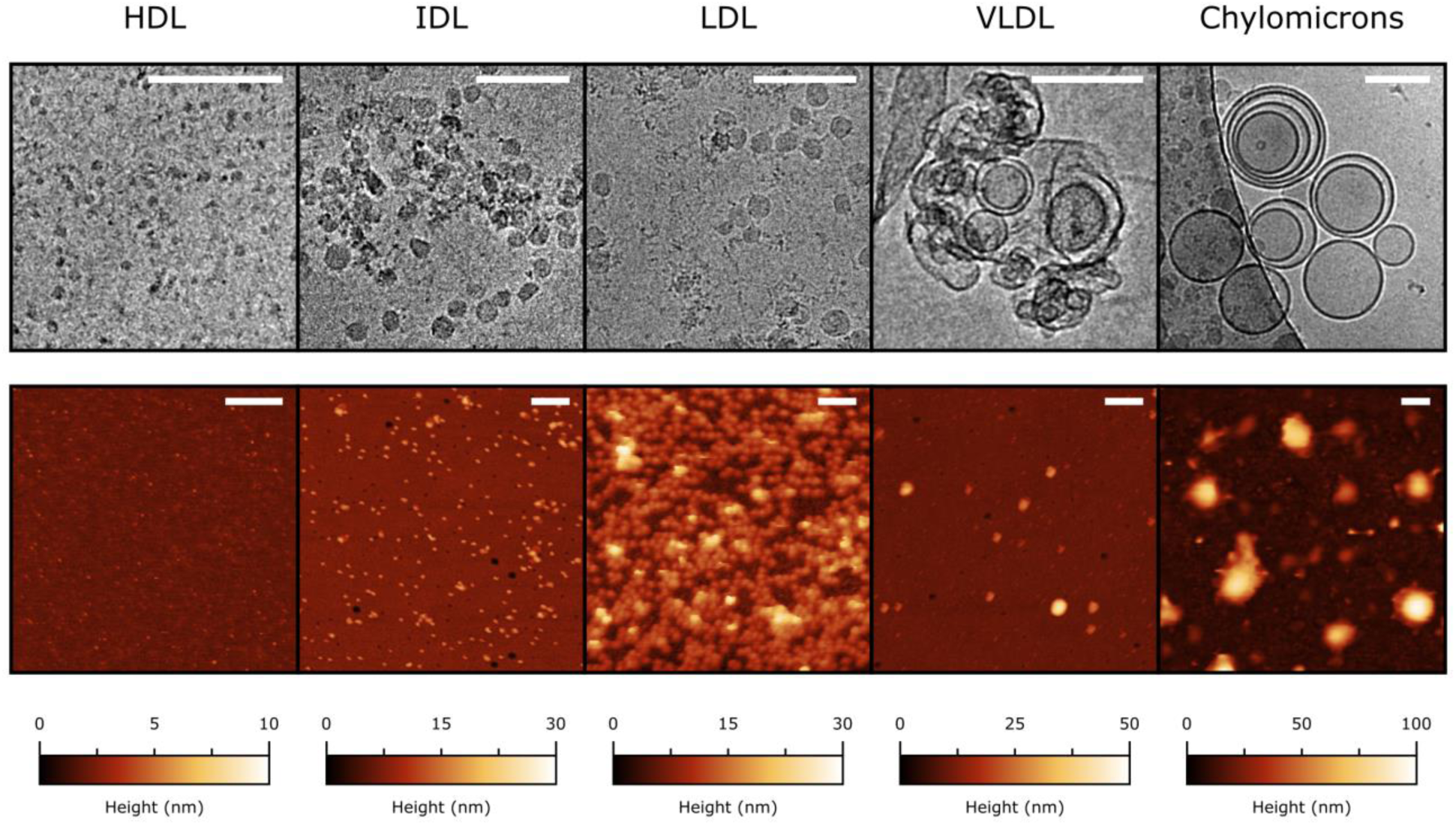
Top: representative cryo-EM images of purified LPs. All cryo-EM scalebars are 100 nm. Compared to EVs, recurring LPs structural features include the lack of decoration and luminal cargo. Chylomicrons and VLDLs show high-contrast boundaries similar in appearance to those displayed by EVs, while LDLs, IDLs and HDLs do not show any discontinuity between bulk and surface. Bottom: representative liquid AFM images of purified LPs. All AFM scalebars are 200 nm.

In particular, EVs appeared as being delimited by a high-contrast boundary decorated with disordered material, most probably corresponding to the lipid bilayer plus associated membrane proteins and glycans. Sporadically, hints of discrete internal cargo could also be detected (Figure 1).

Among LPs, only VLDLs and Chylomicrons were delimited by high-contrast boundaries, while HDLs, IDLs and LDLs did not show significant differences in contrast between their inner and external regions (Figure 2). No sign of external decoration or internal cargo was detectable in any LP sample. A substantial proportion of chylomicrons and VLDLs showed concentric boundaries, which might correspond to the EM projection of either multilamellar or highly corrugated objects. While the univocal attribution of this pattern to a specific structure is challenging, we note that very similar micrographs were previously reported for samples possibly containing these classes of LPs [Baramuikov 2022, Gallart-Palau 2015, Emelyanov 2015]. It is interesting to note how the small size and low contrast of HDLs concur to make them harder to detect and measure in comparison to all other samples, putting them at the practical limit of the technique in this context.

### Single-particle size distributions via cryo-EM and AFM

Cryo-EM and liquid AFM are widely applied to the characterization of vesicle morphology [Robson 2018]. Both techniques can be employed to measure the diameters of several individual vesicles, thus reconstructing their size distribution without resorting to ensemble-averaging. Since EM micrographs can be effectively regarded as two-dimensional projections of the sample [Almgren 2000], and the shape of intact vesicles in solution is essentially spherical, cryo-EM gives direct access to vesicle diameters via simple circular fits of their boundaries (Figure S4). We first analyzed cryo-EM micrographs of all LPs in the series to quantify their size distributions. The largest LPs (Chylomicrons and VLDL) were found to have regular spherical shapes, and their diameters were determined as those of EVs by simple circular fits of their outer boundary. Since the shapes of HDLs, IDLs and LDLs appeared instead to be more irregular (Figure 2), we assigned each object the average of their minimum and maximum Feret diameters (Figure S4, see Materials and Methods).

We then recorded AFM micrographs of all LPs (Figure 2). In contrast to cryo-EM, AFM micrographs cannot convey spherical diameters by direct measuring; however, they contain the three-dimensional profiles of vesicles after deformation due to surface adhesion forces. Most of the recent AFM-based studies agree that vesicles adopt a spherical cap shape upon adsorption on a surface, largely preserving their initial bilayer surface area [Ridolfi 2020a, Vorselen 2017, Vorselen 2020]. Via quantitative AFM morphometry, it is possible to measure the individual surface areas of spherical caps corresponding to each particle in an AFM image, then calculate their equivalent solution diameter (D_eq_), *i*.*e*. the diameter they had in solution in their original spherical shape (Figure S4). Since the validity of this approach was previously demonstrated for vesicles only [Ridolfi 2020a], we checked its applicability to LPs by comparing diameter distributions obtained via cryo-EM and AFM (please refer to the materials and methods section for details) on the whole series of purified LPs (Figure 3).

**Figure 3.**
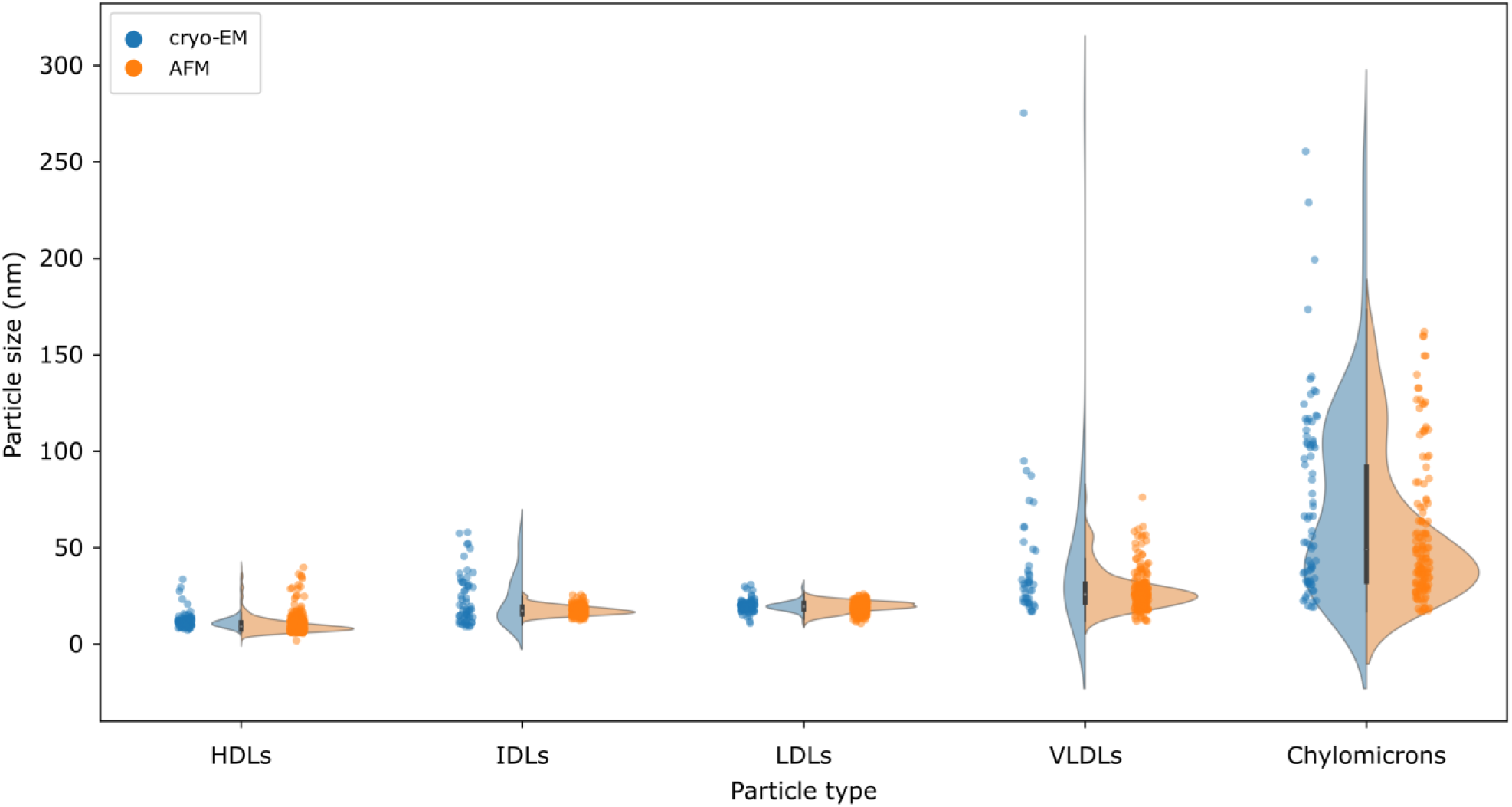
Comparison of size distributions obtained from cryo-EM image analysis (blue dots and frequency plots) and D_eq_ distributions calculated from AFM morphometry (orange dots and frequency plots). The two methods largely agree on the modes of all distributions.

The size distributions reconstructed via both techniques were found to be in very good accord (Figure 3), with nearly coincident main modes and a broad agreement on the position of individual peaks in clearly bimodal distributions (e.g. chylomicrons). Taken together, these measurements show that the same liquid AFM morphometry method developed for EVs [Ridolfi 2020a] can be used to successfully assess size distributions of LPs, and that its results are in very good quantitative accord with cryo-EM.

### Nanomechanics of isolated EVs via AFM morphometry

As previously described [Ridolfi 2020a], AFM morphometry can be used to measure the equivalent spherical cap contact angle (CA) of individual globular objects adhered to the substrate (Figure S4). Collecting the morphometric measurements of several hundred individual objects on a CA versus D_eq_ plot makes it possible to quickly quantify their nanomechanical behaviour.

When the measured object is a pressurized vessel – *e*.*g*. an intact vesicle – its shape upon adsorption on a surface is well approximated by a spherical cap, whose CA is representative of the degree of deformation it experienced during the adhesion, which is in turn determined by its mechanical stiffness [Ridolfi 2021]. Due to this, lipid vesicles will appear as ‘horizontal’ clusters of points on the CA/D_eq_ plot, with a narrow dispersion of CA values across the spanned D_eq_ range. The average CA of the cluster will be proportional to the average stiffness of the objects contributing to form the cluster [Ridolfi 2020a].

Figure 4a contains the CA/D_eq_ plot of EVs enriched and purified from human cardiac progenitor cells cultures (hCPC-EVs)[Andriolo 2018] and of red blood cells-derived EVs (RBC-EVs)[Usman 2018]. The two EV models have been chosen as benchmark EVs derived from different biogenetic pathways and bearing non-identical size distributions. Cryo-EM analysis of hCPC-EVs (see Figure 1) confirmed that intact EVs constituted the vast majority of objects found in this sample. As expected, most Evs in both samples tend to cluster around a characteristic CA value with no dependence on their D_eq_ (Figure 4a), suggesting that these nanoparticles carry the same nanomechanical fingerprint, i.e., similar stiffness independently of their size. Notably, the same result was also previously obtained on different EV populations deposited on substrates identical to those used in this study [Borup 2022]. Due to this, it is possible to define a zone on the CA/D_eq_ plot which is typical of most pressurized EVs. This zone is delimited by 75° < CA < 135°, and D_eq_ > 30 nm.

**Figure 4.**
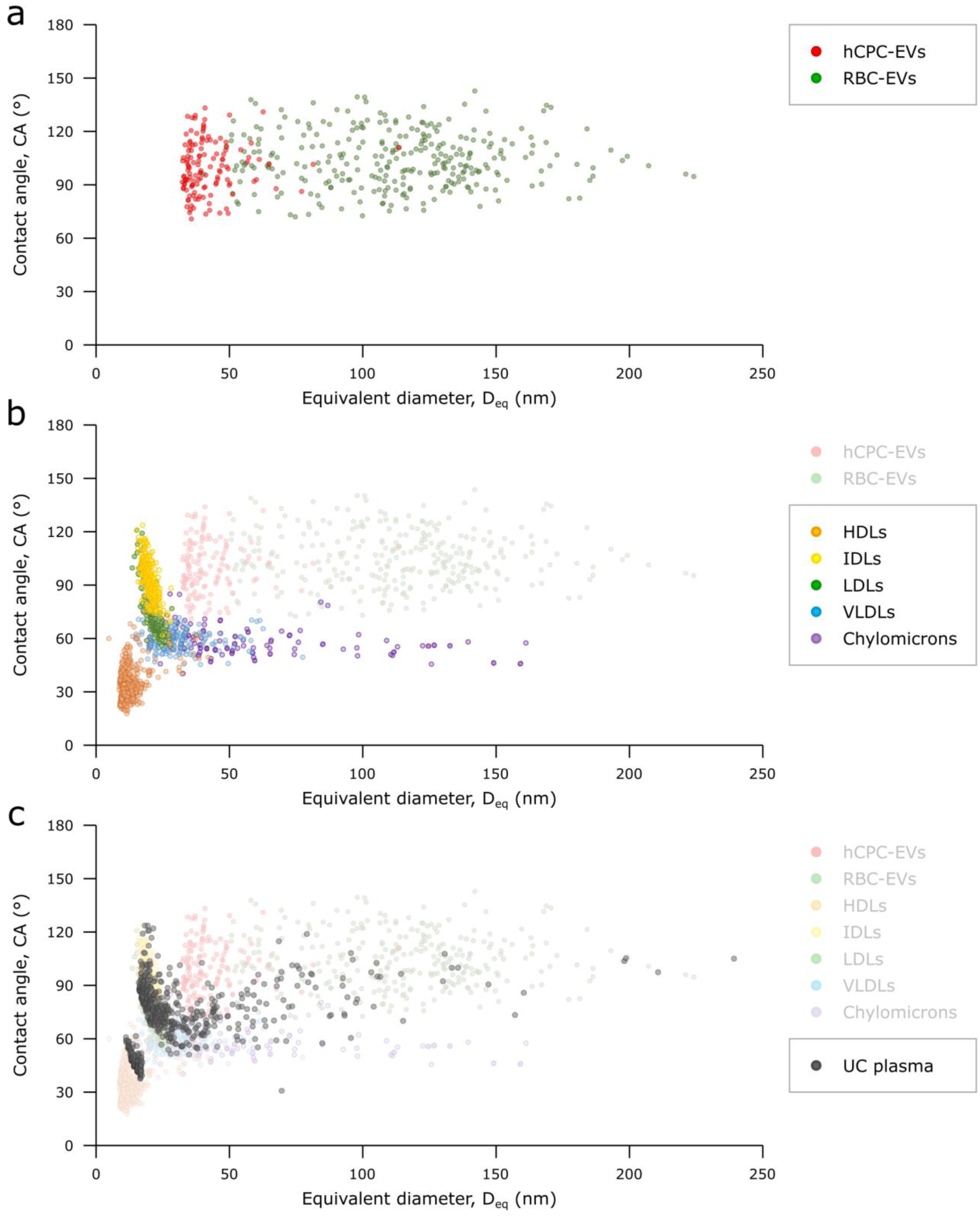
(a): CA/D_eq_ plot of isolated hCPC-EVs (red) and RBC-EVs (green). As expected for objects displaying the nanomechanical behaviour of pressurized elastic vessels, the vast majority of EVs cluster around an average CA value which does not change as a function of D_eq_. In other words, intact EVs have a conserved stiffness which is relatively constant in the small size range associated with EVs. (b): CA/D_eq_ plot of all single-component LP samples: HDL (orange), IDL (yellow), LDL (green), VLDL (light blue), Chylomicrons (purple). The plot in panel (a) is superimposed for the sake of comparison. Each LP subclass populates a different cluster with marginal overlap; no LP subtype populates the zone previously associated with EVs. Chylomicrons and VLDLs are characterized by a conserved average CA value, making their nanomechanical behaviour similar to that of EVs but with lower stiffness. Smaller LPs evidence different (and currently unknown) mechanical behaviours. (c): CA/D_eq_ plot of an ultracentrifuged plasma sample containing a variety of LPs as well as EVs (black). Plots (a) and (b) are superimposed for the sake of comparison. The plot contains the AFM morphometrical measurements of several hundred individual objects found within the plasma sample, making it possible to estimate the presence of specific LP classes, EVs, and their relative abundance (see Figure S5).

### Nanomechanics of single-component LP samples via AFM morphometry

We then applied the same morphometry-based nanomechanical assessment to all LPs in the series. As mentioned above, this approach is in principle only valid for particles characterized by the nanomechanics of pressurized vessels, such as vesicles. However, the fact that independent AFM and cryo-EM morphometry measurements of LPs give results in quantitative agreement [Figure 3] is a strong indication that the morphological parameters used to calculate D_eq_ from AFM images (*i*.*e*. the height and projected diameter of adhered objects [Ridolfi 2020a]) are robust geometrical descriptors for LPs as they were shown to be for EVs. Since CAs are calculated based on the same morphological parameters (Figure S4), our hypothesis is that the position of individual LPs on the same CA/D_eq_ plots used for vesicles could contain an additional layer of information with respect to what could be inferred from their size distributions alone, even in absence of an established model for the nanomechanics of LPs.

Following this approach, we found that different classes of LPs form distinct clusters with marginal overlapping between each other and EVs (Figure 4b). In particular, the clusters of IDLs and LDLs overlap partially, as do those of VLDLs and Chylomicrons; it is uncertain whether this is due to cross-contamination resulting from imperfect separation, or to a high degree of physicochemical similitude between compositionally different populations. Nevertheless, none of the LP clusters populates the zone previously assigned to EVs on the CA/D_eq_ plot, and each LP cluster has a distinctly different centroid even when marginally overlapping other clusters as discussed above.

It is important to note that it would be impossible to distinguish some of the samples based on their size distributions alone. For example, there is a significant overlap in sizes between EVs and larger LPs such as VLDLs and Chylomicrons, as well as between all the smallest LP subclasses. Conversely, combining the CA and D_eq_ values of each nanoparticle enables the differentiation between different LP subtypes and EVs, and gives hints about their nanomechanical behaviour.

In particular, VLDLs and Chylomicrons – despite exhibiting EV-compatible sizes – show a much lower average CA, meaning that their mechanical stiffness is lower than that of intact EVs. Interestingly, they both form horizontal clusters with conserved CAs, which – as discussed above – strongly indicates a ‘vesicle-like’ nanomechanical behaviour. Intriguingly, they also are the only LPs to show high-contrast boundaries in cryo-EM; while the structural details of these boundaries remain unclear, it seems logic to link their presence to the ability to withstand pressurization and thus mechanically act as elastic pressurized vessels.

Conversely, LPs not displaying a high-contrast boundary at cryo-EM do not show a ‘vesicle-like’ horizontal cluster on the CA/D_eq_ plot. As such, they do not have a conserved CA and there is no justification to assume they have a well-defined stiffness value. Due to this, while their CAs are most probably still broadly influenced by their mechanical characteristics, their exact relationship is at the moment still uncharacterized. It is thus best to rationalize the CAs of HDLs, IDLs and LDLs just as quantitative descriptors of their general geometry after adhesion. The usefulness of this approach is that their clusters are indeed distinct from both other LPs and EVs on the CA/D_eq_ plot; in other words, the CAs of small LPs – while not necessarily a descriptor of the particles’ stiffness – still allow discriminating between their mechanical behaviour (Figure 4b).

It is important to note that, in the same way as previously discussed for cryo-EM, the very small size of HDLs makes them a special case for which the applicability of quantitative AFM morphometry must be considered borderline. While probe convolution was found to have a very limited practical impact on the nanomechanical assessment of EVs and liposomes [Ridolfi 2020a], HDLs have sizes comparable with the curvature radii of most commercial probes, leading to a considerably increased convolution weight in their apparent morphology. Due to this, HDLs most probably have artefactually low CAs with respect to other LPs and EVs; nevertheless, by always using the same probes and substrates, we found that their position of the CA/D_eq_ plot was very reproducible across multiple batches and experiments.

### Assessment of a multi-component EVs/LPs mixture via AFM morphometry

Taken together, all the results described in the previous section suggest that individual LP subclasses only span specific combinations of CA and D_eq_ values, which can then be used to define characteristic regions in an CA/D_eq_ plot (Figure S5), in the same way as it was previously done for EVs. The position on an CA/D_eq_ plot of individual unknown objects found in a mixed EV/LP sample will thus enable their classification to one specific class of objects.

As a proof of concept, we applied our AFM nanomechanical imaging assay to several hundred (N=745) individual objects found in a sample known to contain both EVs and LPs. Ultracentrifugation (UC) is an extremely widespread EV enrichment technique [Gudbergsson 2016] which relies on the different sedimentation rates of individual components in a complex mixture. However, due to the considerably higher abundance of LPs compared to EVs in blood, UC unavoidably results in the co-sedimentation of large amounts of LPs, in particular HDLs and LDLs [Simonsen 2017], together with EVs. We thus centrifuged a platelet-free plasma sample at 100K g for 120’, resuspended the pellet and analysed its contents via liquid AFM morphometry.

Individual points in the resulting CA/D_eq_ plot ended up populating multiple different regions, representative of both EVs and LPs (Figure 4c). This result is already sufficient to qualitatively assess the presence of EVs and of each class of LPs in the mixture. In this case, our ultracentrifuged plasma sample contains both EVs and LPs, and the numerically most prominent clusters correspond to those LP subclasses known to be most abundantly coisolated with EVs, *i*.*e*. HDLs and LDLs [Simonsen 2017].

We have previously determined that the surface densities of synthetic liposome solutions deposited on PLL-functionalized substrates correlate with their original concentration in solution [Caselli 2021]. By assuming that the same also holds true for natural EV/LP mixtures, it is possible to use their CA/D_eq_ plots to infer further quantitative information on the mixture’s contents. In particular, it is possible to count individual objects found in each of the zones previously assigned to specific components to obtain a compositional distribution of the mixture. For example, only 8% of the individual objects found in our ultracentrifuged plasma sample was found to have EV-compatible sizes and nanomechanics, while HDLs and LDLs accounted for more than half of the sample’s composition [Figure S5].

Moreover, since AFM morphometry also yields accurate size estimations (see above) of individual objects corresponding to each point in the CA/D_eq_ plot, it is possible to express the relative abundances of components in a mixture in terms of surface area and volume instead of number density. For example, EVs were found to constitute 74% of the total exposed surface area of all objects in the mixed plasma sample, and 94% of their volume, despite their population being only the 8%. Conversely, HDLs, IDLs and LDLs were measured to collectively constitute less than 1% of the volume despite being more than 70% of the individual objects.

As stated in previous sections, HDLs proved to be the most problematic class of LPs to characterise via both cryo-EM and AFM morphometry, even when observed in purified samples. Accordingly, their detection and quantification in complex mixtures proved to be even more challenging. In addition to the technical hurdles inherent in their observation (low EM contrast, small size, significant AFM probe convolution), HDLs in mixed LP/EV samples pose additional challenges due to their relative abundance compared to other components. In our centrifuged plasma sample, they qualitatively appeared to be the most abundant among all classes of objects, including other LPs; a pervasive layer of abundant small objects possibly corresponding to HDLs is often visible in EM micrographs of plasma samples [Nieuwland 2022]. However, quantitative cryo-EM and AFM morphometry measurements both necessitate of micrographs in which the analytes appear as discrete, well-resolved objects. The high relative abundance of HDLs implies that if the mixed sample was diluted up to the point of resolving individual HDLs, other components would become vanishingly rare in micrographs. Conversely, if the sample is analyzed at concentrations at which other components are found with reasonable frequency, HDLs are too crowded to be reliably detected and measured. This is what happened in the CA/D_eq_ plot of our ultracentrifuged plasma sample, in which the HDL cluster is clearly populating only a portion of the zone previously assigned to these LPs (Figures 4b,4c), the one corresponding to larger CAs and sizes. Due to these considerations, it is very likely that the relative amount of HDLs in mixed samples is considerably underestimated.

## Conclusions

We herein demonstrated how the AFM morphometry-based method we previously applied to intact EVs can be seamlessly extended to the nanomechanical assessment of LPs and complex mixed EVs/LPs samples. Pooling hundreds of individual objects on the same CA/D_eq_ plot makes it possible to resolve the nanomechanical properties of co-isolated EVs and LPs, and broadly quantify their abundance, hence providing a very useful tool for quickly assessing the purity of several EV/LP isolates including plasma- and serum-derived preparations.

Moreover, we showed how the AFM morphometry-based measurement of D_eq_ provides LPs size distributions in very good agreement with those obtained by cryo-EM. Having access to realistic size estimates of individual LPs/EVs in a mixture makes it possible to give their relative amounts in terms of surface area and volume in addition to their stoichiometric ratios.

Our AFM-based assay is label-free, single-step and relatively quick to perform. It does not involve preparative steps which could adversely impact the sample integrity such as drying or staining; it can be performed in buffers and cell culture media. Importantly, it can be used to analyze samples which prove very challenging to assess with several established techniques due to ensemble-averaging, low sensibility to small particles, or both. Current quantitative shortcomings of our assay are mostly linked to the presence of HDLs. Further studies might allow to give better estimates via empiric calibration procedures.

To the best of our knowledge, these results represent the first single-particle systematic mechanical investigation of different lipoprotein classes. All of the very few existing nanomechanical studies on LPs [*e*.*g*. Baraimukov 2022, Gan 2015] are based on theoretical frameworks such as Derjaguin-Muller-Toporov or Hertzian contact mechanics, whose applicability to the mechanical behaviour of pressurized vessels is highly questionable. A more thorough mechanical characterization of LPs, while certainly auspicable, would necessitate appropriate mechanical models for each type of LP, which are at the moment not available. In this context, our approach proved to be crude but effective, being able to leverage nanomechanics to discriminate between EVs and LPs which cannot be resolved by size alone.

## Acknowledgements

This work was partially funded from the European Union’s Horizon 2020 research and innovation program under grant agreements No. 952183 (project BOW) and No. 951768 (project MARVEL). This research has also received funding from MIUR through PRIN 2017E3A2NR_004 project. We acknowledge the Florence Center for Electron Nanoscopy (FloCEN) at the University of Florence and the SPM@ISMN facility at CNR Bologna.

## Notes

### Competing Interest Statement

The authors have declared no competing interest.

## Bibliography

Almgren, M., Edwards, K., & Karlsson, G. (2000). Cryo transmission electron microscopy of liposomes and related structures. Colloids and Surfaces A: Physicochemical and Engineering Aspects, 174(1–2), 3–21. https://doi.org/10.1016/S0927-7757(00)00516-1

Andriolo, G., Provasi, E., lo Cicero, V., Brambilla, A., Soncin, S., Torre, T., Milano, G., Biemmi, V., Vassalli, G., Turchetto, L., Barile, L., & Radrizzani, M. (2018). Exosomes from human cardiac progenitor cells for therapeutic applications: Development of a GMP-grade manufacturing method. Frontiers in Physiology, 9(AUG), 1169. https://doi.org/10.3389/FPHYS.2018.01169

Bairamukov, V. Yu., Bukatin, A. S., Kamyshinsky, R. A., Burdakov, V. S., Pichkur, E. B., Shtam, T. A., & Starodubtseva, M. N. (2022). Nanomechanical characterization of exosomes and concomitant nanoparticles from blood plasma by PeakForce AFM in liquid. Biochimica et Biophysica Acta (BBA) - General Subjects, 1866(7), 130139. https://doi.org/10.1016/j.bbagen.2022.130139

Borup, A., Boysen, A. T., Ridolfi, A., Brucale, M., Valle, F., Paolini, L., Bergese, P., & Nejsum, P. (2022). Comparison of separation methods for immunomodulatory extracellular vesicles from helminths. Journal of Extracellular Biology, 1(5). https://doi.org/10.1002/jex2.41

Botha, J., Handberg, A., & Simonsen, J. B. (2022). Lipid-based strategies used to identify extracellular vesicles in flow cytometry can be confounded by lipoproteins: Evaluations of annexin V, lactadherin, and detergent lysis. Journal of Extracellular Vesicles. https://doi.org/10.1002/jev2.12200

Brennan, K., Martin, K., FitzGerald, S. P., O’Sullivan, J., Wu, Y., Blanco, A., Richardson, C., & Mc Gee, M. M. (2020). A comparison of methods for the isolation and separation of extracellular vesicles from protein and lipid particles in human serum. Scientific Reports, 10(1). https://doi.org/10.1038/S41598-020-57497-7

Busatto, S., Zendrini, A., Radeghieri, A., Paolini, L., Romano, M., Presta, M., & Bergese, P. (2019). The nanostructured secretome. Biomaterials Science, 8(1), 39–63. https://doi.org/10.1039/C9BM01007F

Busatto, S., Yang, Y., Walker, S. A., Davidovich, I., Lin, W.-H., Lewis-Tuffin, L., Anastasiadis, P. Z., Sarkaria, J., Talmon, Y., Wurtz, G., & Wolfram, J. (2020). Brain metastases-derived extracellular vesicles induce binding and aggregation of low-density lipoprotein. J Nanobiotechnol, 18, 162. https://doi.org/10.1186/s12951-020-00722-2

Busatto, S., Yang, Y., Iannotta, D., Davidovich, I., Talmon, Y., & Wolfram, J. (2022). Considerations for extracellular vesicle and lipoprotein interactions in cell culture assays. Journal of Extracellular Vescicles. https://doi.org/10.1002/jev2.12202

Caselli, L., Ridolfi, A., Cardellini, J., Sharpnack, L., Paolini, L., Brucale, M., Valle, F., Montis, C., Bergese, P., & Berti, D. (2021). A plasmon-based nanoruler to probe the mechanical properties of synthetic and biogenic nanosized lipid vesicles. Nanoscale Horizons, 6(7), 543–550. https://doi.org/10.1039/d1nh00012h

de Rond, L., Libregts, S. F. W. M., Rikkert, L. G., Hau, C. M., van der Pol, E., Nieuwland, R., van Leeuwen, T. G., & Coumans, F. A. W. (2019). Refractive index to evaluate staining specificity of extracellular vesicles by flow cytometry. https://Doi.Org/10.1080/20013078.2019.1643671, 8(1). https://doi.org/10.1080/20013078.2019.1643671

Emelyanov, A., Shtamid, T., Kamyshinsky, R., Garaeva, L., Verlov, N., Miliukhina, I., Kudrevatykh, A., Gavrilov, G., Zabrodskaya, Y., Pchelina, S., & Konevega, A. (2020). Cryo-electron microscopy of extracellular vesicles from cerebrospinal fluid. PLoS ONE. https://doi.org/10.1371/journal.pone.0227949

Freitas, D., Balmaña, M., Poças, J., Campos, D., Osório, H., Konstantinidi, A., Vakhrushev, S. Y., Magalhães, A., & Reis, C. A. (2019). Different isolation approaches lead to diverse glycosylated extracellular vesicle populations. Journal of Extracellular Vesicles, 8(1). https://doi.org/10.1080/20013078.2019.1621131

Gallart-Palau, X., Serra, A., See, A., Wong, W., Sandin, S., Lai, M. K. P., Chen, C. P., Kon, O. L., & Kwan Sze, S. (2015). Extracellular vesicles are rapidly purified from human plasma by PRotein Organic Solvent PRecipitation (PROSPR). Nature Publishing Group, 5, 14664. https://doi.org/10.1038/srep14664

Gan, C., Ao, M., Liu, Z., & Chen, Y. (2015). Imaging and force measurement of LDL and HDL by AFM in air and liquid. FEBS Open Bio, 5, 276. https://doi.org/10.1016/J.FOB.2015.03.014

Geeurickx, E., & Hendrix, A. (2020). Targets, pitfalls and reference materials for liquid biopsy tests in cancer diagnostics. Molecular Aspects of Medicine, 72, 100828. https://doi.org/10.1016/J.MAM.2019.10.005

Gudbergsson, J. M., Johnsen, K. B., Skov, M. N., & Duroux, M. (2016). Systematic review of factors influencing extracellular vesicle yield from cell cultures. Cytotechnology, 68(4), 579. https://doi.org/10.1007/S10616-015-9913-6

Holcar, M., Kandušer, M., & Lenassi, M. (2021). Blood Nanoparticles - Influence on Extracellular Vesicle Isolation and Characterization. Frontiers in Pharmacology, 12. https://doi.org/10.3389/FPHAR.2021.773844

Hu, Y., Thaler, J., & Nieuwland, R. (2021). Extracellular Vesicles in Human Milk. Pharmaceuticals (Basel, Switzerland), 14(10). https://doi.org/10.3390/PH14101050

Hutter, J. L., & Bechhoefer, J. (1993). Calibration of atomic-force microscope tips. Review of Scientific Instruments, 64(7), 1868–1873. https://doi.org/10.1063/1.1143970

Johnsen, K. B., Gudbergsson, J. M., Andresen, T. L., & Simonsen, J. B. (2019). What is the blood concentration of extracellular vesicles? Implications for the use of extracellular vesicles as blood-borne biomarkers of cancer. Biochimica et Biophysica Acta - Reviews on Cancer, 1871(1), 109–116. https://doi.org/10.1016/J.BBCAN.2018.11.006

LeClaire, M., Gimzewski, J., & Sharma, S. (2021). A review of the biomechanical properties of single extracellular vesicles. Nano Select, 2(1), 1–15. https://doi.org/10.1002/NANO.202000129

Maas, S. L. N., Breakefield, X. O., & Weaver, A. M. (2017). Extracellular Vesicles: Unique Intercellular Delivery Vehicles. Trends in Cell Biology, 27(3), 172–188. https://doi.org/10.1016/J.TCB.2016.11.003

Mørk, M., Handberga, A., Pedersen, S., Jørgensen, M. M., Bæk, R., Nielsen, M. K., & Kristensen, S. R. (2017). Prospects and limitations of antibody-mediated clearing of lipoproteins from blood plasma prior to nanoparticle tracking analysis of extracellular vesicles. Journal of Extracellular Vesicles, 6(1). https://doi.org/10.1080/20013078.2017.1308779

Necas, D., & Klapetek, P. (2012). Gwyddion: An open-source software for SPM data analysis. Central European Journal of Physics, 10(1), 181–188. https://doi.org/10.2478/s11534-011-0096-2

Nieuwland, R., Siljander, P. R.-M., Falcón-Pérez, J. M., & Witwer, K. W. (2022). Reproducibility of extracellular vesicle research. European Journal of Cell Biology, 101(3), 151226. https://doi.org/10.1016/J.EJCB.2022.151226

Piontek, M. C., Lira, R. B., & Roos, W. H. (2021). Active probing of the mechanical properties of biological and synthetic vesicles. Biochimica et Biophysica Acta. General Subjects, 1865(4). https://doi.org/10.1016/J.BBAGEN.2019.129486

Ramasamy, I. (2014). Recent advances in physiological lipoprotein metabolism. In Clinical Chemistry and Laboratory Medicine (Vol. 52, Issue 12, pp. 1695–1727). Walter de Gruyter GmbH. https://doi.org/10.1515/cclm-2013-0358

Ridolfi, A., Brucale, M., Montis, C., Caselli, L., Paolini, L., Borup, A., Boysen, A. T., Loria, F., van Herwijnen, M. J. C., Kleinjan, M., Nejsum, P., Zarovni, N., Wauben, M. H. M., Berti, D., Bergese, P., & Valle, F. (2020). AFM-Based High-Throughput Nanomechanical Screening of Single Extracellular Vesicles. Analytical Chemistry, 92(15), 10274–10282. https://doi.org/10.1021/acs.analchem.9b05716

Ridolfi, A., Caselli, L., Montis, C., Mangiapia, G., Berti, D., Brucale, M., & Valle, F. (2020). Gold nanoparticles interacting with synthetic lipid rafts: an AFM investigation. Journal of Microscopy, 280(3), 194–203. https://doi.org/10.1111/jmi.12910

Ridolfi, A., Caselli, L., Baldoni, M., Montis, C., Mercuri, F., Berti, D., Valle, F., & Brucale, M. (2021). Stiffness of Fluid and Gel Phase Lipid Nanovesicles: Weighting the Contributions of Membrane Bending Modulus and Luminal Pressurization. Langmuir, 37(41), 12027–12037. https://doi.org/10.1021/acs.langmuir.1c01660

Robson, A. L., Dastoor, P. C., Flynn, J., Palmer, W., Martin, A., Smith, D. W., Woldu, A., & Hua, S. (2018). Advantages and limitations of current imaging techniques for characterizing liposome morphology. Frontiers in Pharmacology, 9(FEB), 80. https://doi.org/10.3389/fphar.2018.00080

Schindelin, J., Arganda-Carreras, I., Frise, E., Kaynig, V., Longair, M., Pietzsch, T., Preibisch, S., Rueden, C., Saalfeld, S., Schmid, B., Tinevez, J. Y., White, D. J., Hartenstein, V., Eliceiri, K., Tomancak, P., & Cardona, A. (2012). Fiji: an open-source platform for biological-image analysis. Nature Methods 2012 9:7, 9(7), 676–682. https://doi.org/10.1038/nmeth.2019

Simonsen, J. B. (2017). What are we looking at? Extracellular vesicles, lipoproteins, or both? In Circulation Research (Vol. 121, Issue 8, pp. 920–922). Lippincott Williams and Wilkins. https://doi.org/10.1161/CIRCRESAHA.117.311767

Sloop, C. H., Dory, L., & Roheim, P. S. (1987). Interstitial fluid lipoproteins. Journal of Lipid Research, 28(3), 225–237. https://doi.org/10.1016/S0022-2275(20)38701-0

Sódar, B. W., Kittel, Á., Pálóczi, K., Vukman, K. v., Osteikoetxea, X., Szabó-Taylor, K., Németh, A., Sperlágh, B., Baranyai, T., Giricz, Z., Wiener, Z., Turiák, L., Drahos, L., Pállinger, É., Vékey, K., Ferdinandy, P., Falus, A., & Buzás, E. I. (2016). Low-density lipoprotein mimics blood plasma-derived exosomes and microvesicles during isolation and detection. Scientific Reports 2016 6:1, 6(1), 1–12. https://doi.org/10.1038/srep24316

Sorkin, R., Huisjes, R., Bošković, F., Vorselen, D., Pignatelli, S., Ofir-Birin, Y., Freitas Leal, J. K., Schiller, J., Mullick, D., Roos, W. H., Bosman, G., Regev-Rudzki, N., Schiffelers, R. M., & Wuite, G. J. L. (2018). Nanomechanics of Extracellular Vesicles Reveals Vesiculation Pathways. Small, 14(39), 1801650. https://doi.org/10.1002/SMLL.201801650

Usman, W. M., Pham, T. C., Kwok, Y. Y., Vu, L. T., Ma, V., Peng, B., Chan, Y. S., Wei, L., Chin, S. M., Azad, A., He, A. B. L., Leung, A. Y. H., Yang, M., Shyh-Chang, N., Cho, W. C., Shi, J., & Le, M. T. N. (2018). Efficient RNA drug delivery using red blood cell extracellular vesicles. Nature Communications 2018 9:1, 9(1), 1–15. https://doi.org/10.1038/s41467-018-04791-8

van der Pol, E., de Rond, L., Coumans, F. A. W., Gool, E. L., Böing, A. N., Sturk, A., Nieuwland, R., & van Leeuwen, T. G. (2018). Absolute sizing and label-free identification of extracellular vesicles by flow cytometry. Nanomedicine : Nanotechnology, Biology, and Medicine, 14(3), 801–810. https://doi.org/10.1016/J.NANO.2017.12.012

Vorselen, D., Mackintosh, F. C., Roos, W. H., & Wuite, G. J. L. (2017). Competition between Bending and Internal Pressure Governs the Mechanics of Fluid Nanovesicles. ACS Nano, 11(3), 2628–2636. https://pubs.acs.org/doi/10.1021/acsnano.6b07302

Vorselen, D., van Dommelen, S. M., Sorkin, R., Piontek, M. C., Schiller, J., Döpp, S. T., Kooijmans, S. A. A., van Oirschot, B. A., Versluijs, B. A., Bierings, M. B., van Wijk, R., Schiffelers, R. M., Wuite, G. J. L., & Roos, W. H. (2018). The fluid membrane determines mechanics of erythrocyte extracellular vesicles and is softened in hereditary spherocytosis. Nature Communications 2018 9:1, 9(1), 1–9. https://doi.org/10.1038/s41467-018-07445-x

Vorselen, D., Piontek, M. C., Roos, W. H., & Wuite, G. J. L. (2020). Mechanical Characterization of Liposomes and Extracellular Vesicles, a Protocol. Frontiers in Molecular Biosciences, 7, 139. https://doi.org/10.3389/FMOLB.2020.00139/BIBTEX

Zhang, X., Borg, E. G. F., Liaci, A. M., Vos, H. R., & Stoorvogel, W. (2020). A novel three step protocol to isolate extracellular vesicles from plasma or cell culture medium with both high yield and purity. Journal of Extracellular Vesicles, 9(1), 1791450. https://doi.org/10.1080/20013078.2020.1791450

