## Supporting Information for "Compositional profiling of EV-lipoprotein mixtures by AFM nanomechanical imaging"

**Preparation of RBC-EVs**

Blood samples were obtained by healthy donors from Spedali Civili hospital (Brescia, Italy) with informed consents. All experiments with human blood samples were performed according to the guidelines and the approval of the Spedali Civili Ethics committee.

RBC-EVs were isolated following the protocol from Usman et al [Usman 2018, Nyugen 2016]. Briefly, after blood collection, RBCs were pelleted by centrifugation at 1000 × *g* for 8 minutes at 4°C (5804R Eppendorf centrifuge, A-4-44 rotor, 15 ml tubes) and washed three times in PBS (Corning, USA). RBCs were further washed two times with CPBS [PBS + 0.1 g/L calcium chloride (Sigma Aldrich, St. Louis, USA)] and transferred into a 75 $mm^2$ tissue culture flask (Corning, USA). Calcium ionophore (A23187, Sigma Aldrich, St. Louis, USA) was added to the flask (final concentration 10 mM) and incubated overnight at 37°C.

To separate EVs, RBCs and cell debris were removed by differential centrifugation (600 × *g* for 20 min, 1600 × *g* for 15 min, 3260 × *g* for 15 min, and 10,000 × *g* for 30 min at 4 °C). The pellet was discarded at every step, transferring the supernatant to a fresh tube. The supernatants were filtered through 0.45 µm nylon syringe filters (Sarstedt, Germany).

EVs were concentrated by ultracentrifugation at 100,000 × g for 70 min at 4 °C (Optima XPN-100, TY45 Ti rotor, Beckman Coulter, USA). EV pellets were then resuspended in cold PBS, layered above 2 ml frozen 60% sucrose cushion and centrifuged at 100,000 × *g* for 16 h at 4 °C (Optima MAX-XP, MLS-50 rotor, Beckman Coulter, USA), with deceleration speed set to 0. The red layer of EVs was collected and washed twice with cold PBS and spun at 100,000 × *g* for 70 min at 4 °C (Optima MAX-XP, TLA-55 rotor, Beckman Coulter). Finally, EVs were resuspended in 1 ml of cold PBS.

[Nguyen 2016]: Nguyen, D. B., Thuy Ly, T. B., Wesseling, M. C., Hittinger, M., Torge, A., Devitt, A., Perrie, Y., & Bernhardt, I. (2016). Characterization of Microvesicles Released from Human Red Blood Cells. Cellular Physiology and Biochemistry : International Journal of Experimental Cellular Physiology, Biochemistry, and Pharmacology, 38(3), 1085–1099. https://doi.org/10.1159/000443059

[Usman 2018]: Usman, W. M., Pham, T. C., Kwok, Y. Y., Vu, L. T., Ma, V., Peng, B., Chan, Y. S., Wei, L., Chin, S. M., Azad, A., He, A. B. L., Leung, A. Y. H., Yang, M., Shyh-Chang, N., Cho, W. C., Shi, J., & Le, M. T. N. (2018). Efficient RNA drug delivery using red blood cell extracellular vesicles. Nature Communications 2018 9:1, 9(1), 1–15. https://doi.org/10.1038/s41467-018-04791-8

**Preparation of hCPC EVs**

EVs were isolated from media conditioned (CM) by CPC . Briefly, CPCs (70-80 % confluent) were washed twice with DPBS and incubated at 37°C with 5% $CO_2$ in DMEM 4.5 g/l glucose without phenol red (Gibco/Thermo

Fisher Scientific). After 7 days, the CM was clarified by 0.22 μm filtration through bottle filter units or on-line filters (ULTA Capsule HC, KMP-HC9202HH, GE Healthcare, USA). Concentration and EV size selection were performed by tangential flow filtration (TFF), using the ÄKTA™ flux 6 system (GE Healthcare) equipped with a 300 kDa cut-off hollow fiber cartridge (GE Healthcare); the concentration was followed by diafiltration in 5 volumes of Plasma-Lyte A® solution.

| ***Sample*** | ***Particle/mL (NTA)*** | ***Diameter (NTA)*** | Diameter (***DLS)*** |
|---|---|---|---|
| *HDL* | *ND* | *ND* | *10.4 ± 0.4 nm* |
| *LDL* | *ND* | *ND* | *17.4 ± 2.9 nm* |
| *IDL* | *ND* | *ND* | *46.5 ± 3.2 nm* |
| *VLDL* | $2.5*10^{12} \pm 1.2*10^{7}$ | *148 ± 3 nm* | *127.7 ± 1.7 nm* |
| *Chylomicrons* | $3.3*10^{12} \pm 2.3*10^{7}$ | *178 ± 4 nm* | *174.4 ± 4.6 nm* |
| *hCPC-EVs* | $1.0*10^{11} \pm 4.3*10^{6}$ | 193 ± 2 nm | *62.4 ± 1.5 nm* |
| *RBC-EVs* | $1.81*10^{12} \pm 8.8*10^{10}$ | *160 ± 2.3 nm* | *180 ± 4,3 nm* |

***Table S1*** *– determination of particle concentration and diameter by NTA; Hydrodinamic diameter by DLS for lipoproteins, hCPC-EVs, and RBC-EVs samples.*

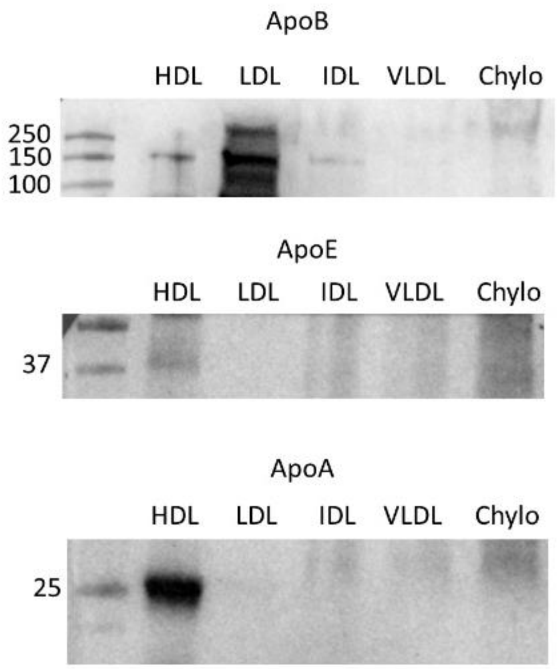


***Figure S1*** *– Western Blotting analysis for Lipoprotein samples confirms the presence of ApoB (enriched in LDL), ApoE and ApoA (enriched in HDL)*

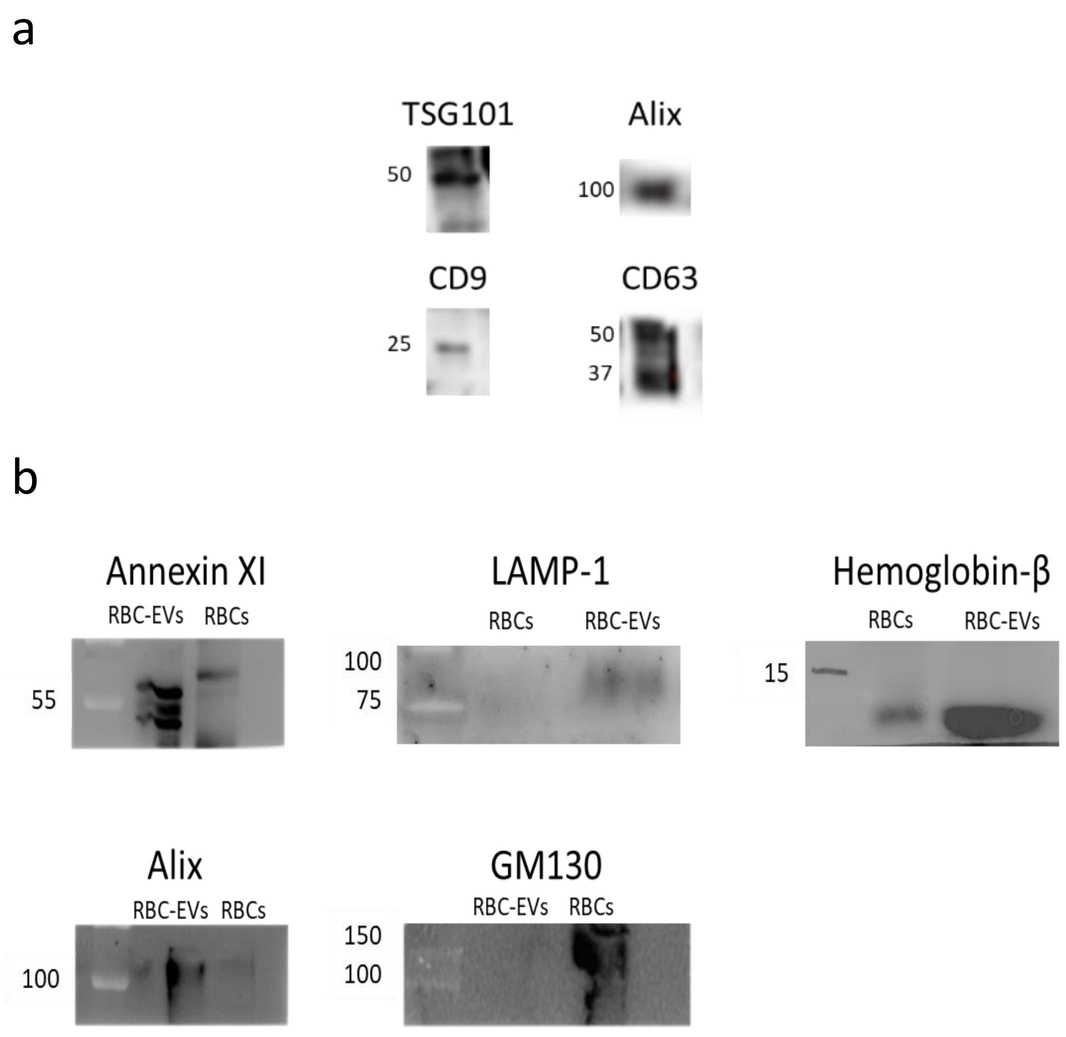


***Figure S2*** *– (a): Western Blotting analysis for the hCPC EVs samples confirms the presence of luminal proteins TSG101 and Alix and of transmembrane proteins CD9 and CD63.*
*(b): Western Blotting analysis for the RBC-EVs samples confirms the presence of luminal proteins Hemoglobin-β and Alix and of transmembrane and membrane-associated proteins LAMP-1 and Annexin XI. Negligible Golgi contaminant GM130 is present in the preparation.*

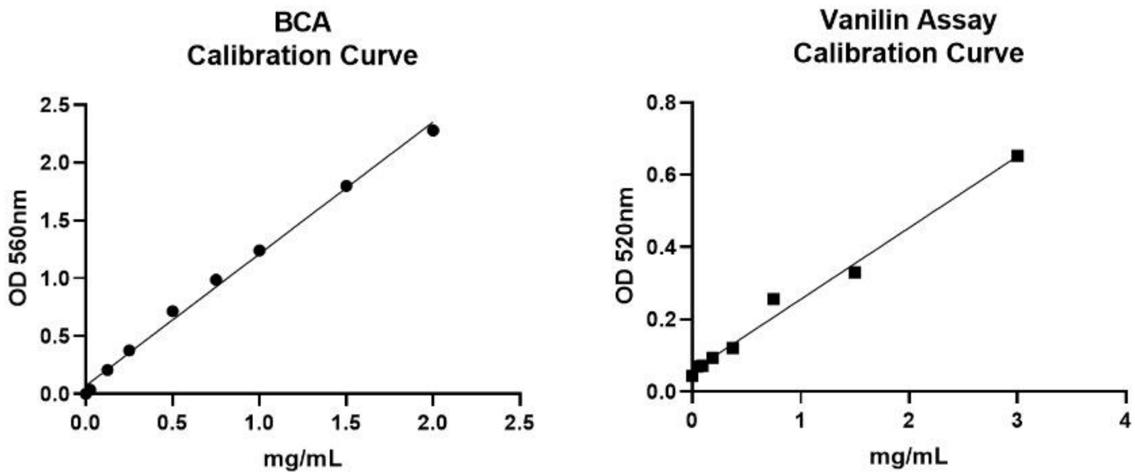


| Sample | BCA (mg/mL) | Lipids (mg/mL) | Ratio |
|---|---|---|---|
| HDL | 402,21 | 285,69 | 1,41 |
| LDL | 347,64 | 462,01 | 0,75 |
| IDL | 0,47 | 2,61 | 0,18 |
| VLDL | 0,45 | 1,75 | 0,26 |
| Chilo | 1,37 | 15,30 | 0,09 |
| EVs | 2,10 | 1,49 | 1,40 |

***Figure S3*** *– BCA and Lipid concentration assay for Lipoproteins and hCPC EVs. Upper panels: calibration curves run with Bovine Serum Albumin (BSA) for the BCA test on the left and DOPC standards for* sulfo-phopsho-vanillin *test on the right. Accordingly, table reports protein content, lipid content and protein/lipid ratio for each analyzed sample.*

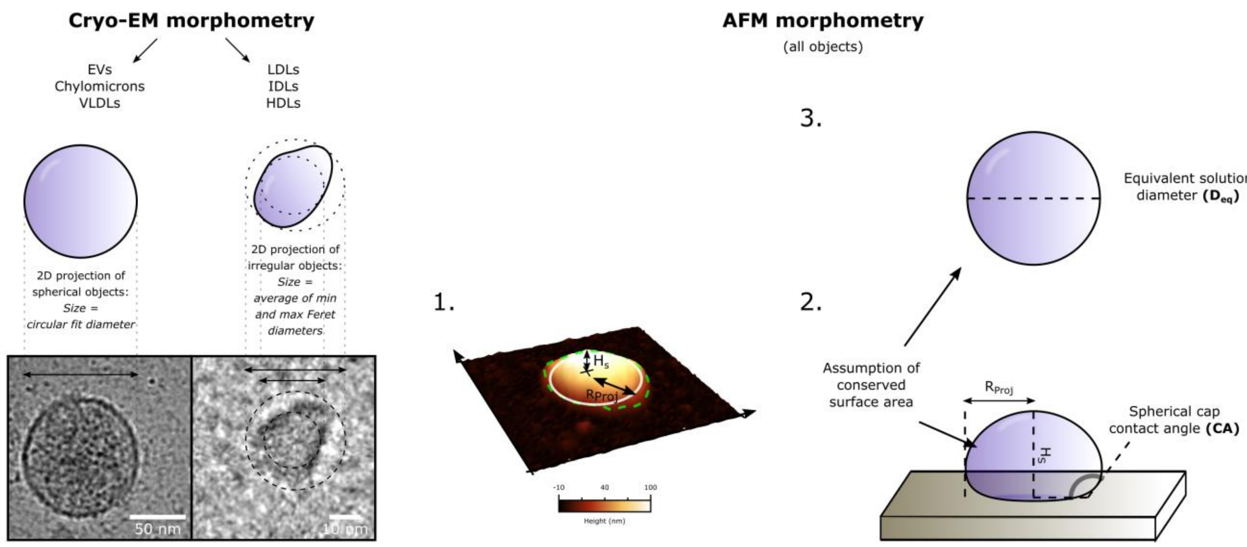


***Figure S4*** *– (Left): Quantitative Cryo-EM morphometry procedure for spherical (EVs, Chylomicrons, VLDLs) and irregular (LDLs, IDLS, HDLs) objects. Spherical objects: direct circular fit. Irregular objects: average of maximum and minimum Feret diameters (see materials and methods). The resulting size distributions are shown in main text figure 3.*

*(Right): Measurement of the equivalent solution diameter ($D_{eq}$) and equivalent spherical cap contact angle (CA) of an unknown particle via AFM morphometry. (1): The height ($H_s$) and maximum inscribed disc radius ($R_{proj}$) of discrete objects are measured on AFM micrographs. (2): The object's geometry is approximated with a spherical cap of height $H_s$ and projected radius $R_{proj.}$ It is then possible to calculate its surface contact angle (CA). Please refer to main text reference [Ridolfi 2020a] for additional details on the procedure. (3): The diameter of a sphere having the same surface area of the spherical cap introduced at step 2 is calculated. The resulting 'equivalent solution siameter' ($D_{eq}$) size distributions are shown in figure 3. It is unnecessary to know whether the analysed particle is an EV or a LP. However, if the particle's nanomechanics are those of an intact vesicle, the CA recapitulates its mechanical stiffness (see main text); if the particle's nanomechanical behaviour is not known (as for LPs), CA is just a robust numerical descriptor of their geometry. CA values from hundreds of individual objects were pooled in the CA/Deq plots shown in main text figure 4.*

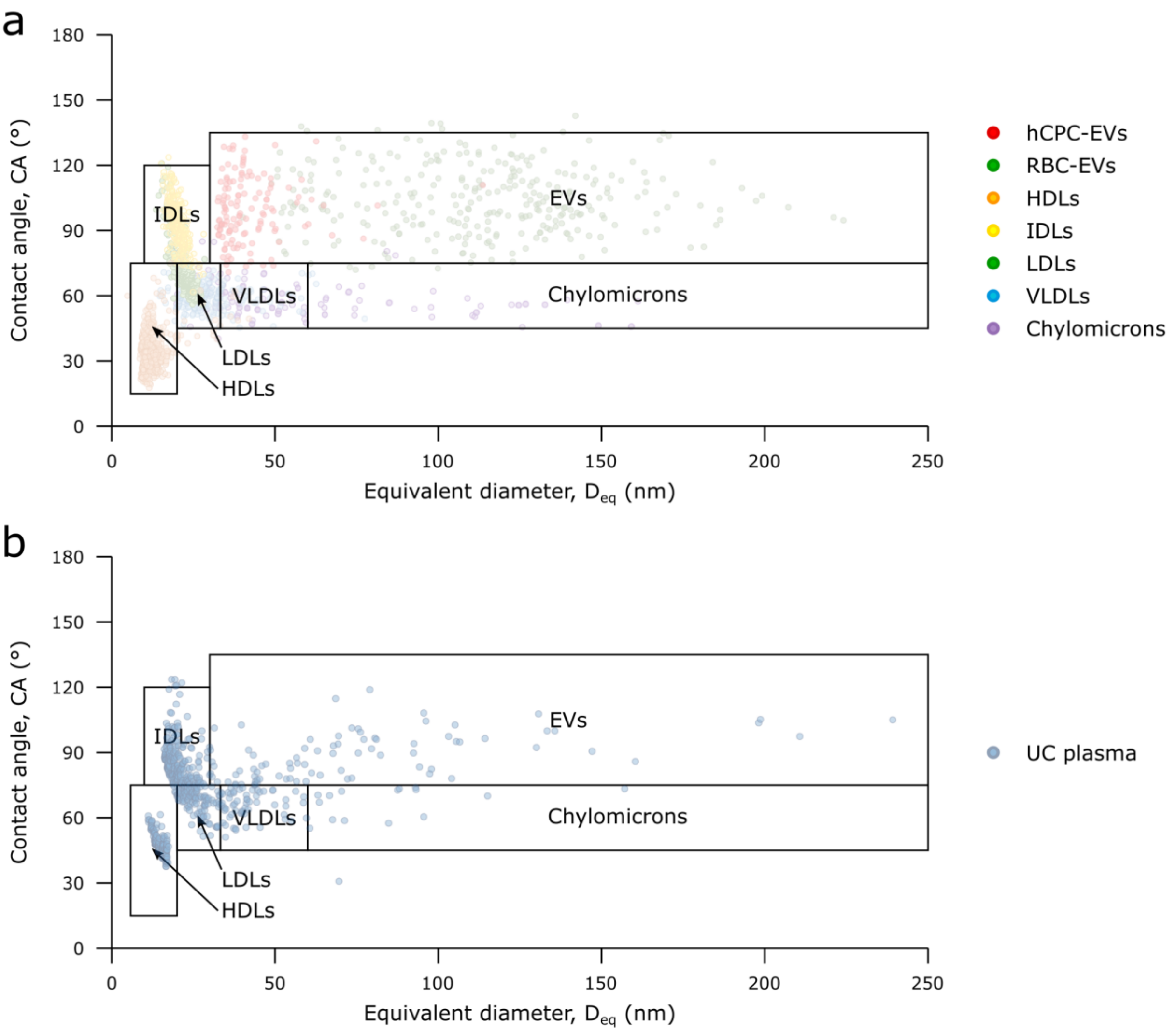


| Relative abundances of ultracentrifuged plasma sample (see main text Figure 4) | | | |
|---|---|---|---|
| *Particle Type* | *Number %* | *Surface area %* | *Volume %* |
| EVs | 8 % | 74 % | 94.8 % |
| Chylomicrons | 3 % | 9 % | 3.2 % |
| VLDLs | 16 % | 9 % | 1.5 % |
| LDLs | 19 % | 3 % | 0.3 % |
| IDLs | 22 % | 3 % | 0.2 % |
| HDLs | 32*% | 2*% | 0.1*% |

* relative amount of HDLs probably underestimated (see main text)

***Figure S5*** *– (a): assignment of biological identity to discrete zones of a CA/$D_{eq}$ scatterplot. (b): Relative abundances in the ultracentrifuged plasma sample (see main text Figure 4c) as estimated via the assignments shown in panel (a).*
